# A common human brain-derived neurotrophic factor polymorphism leads to sustained depression of excitatory synaptic transmission by isoflurane in hippocampus

**DOI:** 10.1101/2021.12.30.474566

**Authors:** RA Williams, KW Johnson, FS Lee, HC Hemmings, J Platholi

## Abstract

Multiple presynaptic and postsynaptic targets have been identified for the reversible neurophysiological effects of general anesthetics on synaptic transmission and neuronal excitability. However, the synaptic mechanisms involved in persistent depression of synaptic transmission resulting in more prolonged neurological dysfunction following anesthesia are less clear. Here, we show that brain-derived neurotrophic factor (BDNF), a growth factor implicated in synaptic plasticity and dysfunction, enhances glutamate synaptic vesicle exocytosis, and that attenuation of vesicular BDNF release by isoflurane contributes to transient depression of excitatory synaptic transmission in mice. This reduction in synaptic vesicle exocytosis was irreversible in neurons that release less endogenous BDNF due to a polymorphism (BDNF Val66Met) compared to wild-type mouse hippocampal neurons following isoflurane exposure. These effects were prevented by exogenous application of BDNF. Our findings identify a role for a common human BDNF single nucleotide polymorphism (Val66Met; rs6265) in persistent changes of synaptic function following isoflurane exposure. These persistent alterations in excitatory synaptic transmission have important implications for the role of genotype in anesthetic effects on synaptic plasticity and neurocognitive function.

## Introduction

General anesthetics are distinguished by route of administration (inhalation [volatile]-VA or intravenous-IVA), and their diverse chemical structures, molecular targets, and binding sites. Although mechanistically distinct, all general anesthetics alter synaptic transmission by acting both presynaptically and postsynaptically [1, 2]. While these effects are anticipated to be fully reversible upon drug elimination, concerns that exposure to general anesthesia in vulnerable populations can induce lasting impairments in synaptic plasticity and cognitive function remain. Perioperative neurocognitive disorder (PND; [3]) affect up to 41% of patients older than 60 years [4-6] and additional risk factors include pre-existing cognitive impairment or disease [7] as well as education level and socioeconomic status [8]. Several reports also show a possible association between a specific genotype and PND; polymorphisms of the human gene C-reactive protein [9], P-selectin [10], and platelet glycoprotein IIIa [11] suggest an additional vulnerability in genotype may increase susceptibility to PND. Thus, identifying and addressing the role of genetic variation in preoperative risk assessment and recovery from anesthesia will help facilitate individualized mechanism-based use of specific anesthetic agents based on patient factors.

Brain-derived neurotrophic factor (BDNF) is synthesized as a precursor (proBDNF) and is cleaved to its mature form (mBDNF) and released to mediate divergent actions on neuronal survival, neuron structure, and synaptic plasticity [12, 13]. In the adult brain, mBDNF regulates density and morphology of dendritic spines [14, 15], promoting neuronal survival and enhancing synaptic plasticity [16-18]. Studies support a role for down-regulation of mBDNF by anesthetics; in humans, both intravenous (propofol) and volatile (isoflurane) anesthetics reduce plasma BDNF concentration intraoperatively and 24 hr after surgery [19], while epigenetic enhancement of BDNF signaling improves cognitive impairment induced by isoflurane anesthesia in aged rats [20].

Human carriers of the common BDNF Val66Met single nucleotide polymorphism (SNP; rs62645; 30% prevalence [21]) have reduced hippocampal volume, altered synaptic plasticity, and impaired hippocampus-dependent learning and memory [22]. This SNP results in abnormal intracellular trafficking of BDNF leading to lower regulated secretion of mBDNF [23-25]. This allele is also associated with reduced preoperative cognitive function leading to attenuated cortical activity during surgery under volatile anesthesia suggestive of further cognitive decline and susceptibility to PND [8, 26, 27]. Understanding the pharmacogenetic interaction between this common SNP and general anesthesia should provide novel insights into the neurocognitive effects of anesthetics, but the cellular and molecular mechanisms behind the contribution of BDNF and the Val66Met polymorphism to synaptic deficits following general anesthesia have not been explored.

Mechanisms for acute depression of excitatory synaptic transmission by general anesthesia include reduction of neuronal excitability [28] or action potential conduction [29-32], inhibition of Ca^2+^ influx [33, 34] and synaptic vesicle (SV) exocytosis [35-37], and/or blockade of postsynaptic glutamate receptors [38]. Accordingly, our previous work has shown differential reduction of evoked release of various major CNS neurotransmitters by volatile anesthetics [39-41]. For example, isoflurane more potently inhibits glutamatergic compared to GABAergic or dopaminergic action-potential (AP)-evoked synaptic vesicle exocytosis in dissociated primary neurons [33, 37]. These observations are similar to findings in cortical GABAergic neurons *in vivo* [42, 43].

Neurotransmitter release is tightly coupled to Ca^2+^ entering the presynaptic bouton [44]. Various anesthetics differentially act on presynaptic Ca^2+^ signaling via ion channels or vesicle fusion mechanisms to inhibit exocytosis [39, 45] by attenuating release probability and number of functional release sites [46, 47]. Candidate targets include Na^+^, Ca^+^, and K^+^ channels [48-50], along with neurotransmitter transporters [35, 51] and SNARE exocytotic proteins [52-54]. Yet these presynaptic sites of action do not fully explain anesthetic-induced effects on synaptic transmission.

Activity-dependent mBDNF secretion enhances neurotransmitter release by multiple mechanisms that increase intracellular Ca^2+^ [55], directly act on vesicle machinery [56, 57], and/or increase neurotransmitter replenishment and release [58, 59]. Mutation or selective deletion of BDNF results in fewer docked vesicles [60], reduced release probability [61], and reduced expression of synaptic vesicle proteins, such as synaptobrevin and synaptophysin, that regulate neurotransmitter release [60]. These observations suggest that regulation of neurotransmitter release by mBDNF may be a novel presynaptic anesthetic mechanism. Therefore, we tested the hypothesis that BDNF signaling contributes of isoflurane-induced depression of excitatory synaptic transmission and that these effects are enhanced in neurons with reduced BDNF secretion.

## Methods

This study was performed in strict accordance with the recommendations in the Guide for the Care and Use of Laboratory Animals of the National Institutes of Health. The Institutional Animal Care and Use Committee (IACUC) of Weill Cornell Medicine specifically approved this study under protocol #2015-0023. All animals were handled according to this approved protocol, and followed ARRIVE guidelines. All surgical procedures were terminal and anesthesia with isoflurane was used to prevent animal suffering.

### Reagents

Isoflurane was obtained from Abbott (Abbott Park, IL) and all other reagents were sourced as indicated. Rat proBDNF-pHluorin was from Ed Chapman (University of Wisconsin, Madison, WI), and rat vGlut1-pHluorin was from Timothy Ryan (Weill Cornell Medicine, New York, NY).

### Hippocampal neuron culture and transfection

Rat and mouse hippocampal cells were cultured according to Calabrese and Halpain [62]. Briefly, whole hippocampi were dissected from embryonic day 16-18 (E16-18) Sprague Dawley rat or mouse embryos, and the cells dissociated with papain, cultured on 25 mm glass coverslips (Carolina Biological, Burlington, NC) in 35-mm dishes (Corning, Durham, NC) at a density of 100,000-150,000 cells/mm^2^, and maintained in Neurobasal medium (GIBCO, Grand Island, NY) supplemented with SM1 (Stem Cell Technologies, Vancouver, BC, Canada) and 0.5 mM L-glutamine (Sigma-Aldrich, St. Louis, MO). For mouse cultures, heterozygous (Val66Met) male and female mice were paired to yield Val66Val (wt), Val66Met, and Met66Met genotypes. Individual mouse pups were genotyped and cultured separately. Cultures were transfected at 7 days *in vitro* (DIV) by calcium phosphate precipitation with 6-10 μg mCherry cDNA to allow visualization of neuronal morphology and with BDNF-pHluorin or vGlut1-pHluorin cDNA to visualize exocytosis as described [63]. Cells were incubated with the transfection mixture for 2.5-3 h in 95% air/5% CO_2_ at 37°, washed twice with pre-warmed HBS solution (in mM: 135 NaCl, 4 KCl, 1 Na_2_HPO_2_, 2 CaCl_2_, 1 MgCl_2_, 10 glucose, and 20 HEPES [pH 7.35]), and replaced with Neurobasal medium. Cells were used for live cell-imaging at 14-16 DIV.

### Drug treatments

Experiments used isoflurane at aqueous concentrations equivalent to 0.25-2 times minimum alveolar concentration (MAC) in rat [64] as a clinically relevant dose range (1 MAC is ED_50_). Saturated stock solutions were diluted to working concentrations for superfusion through Teflon tubing using gas-tight glass syringes. Coverslips were placed in a heated (37°C) perfusion chamber (260 μl) and Tyrode’s solution (in mM: 119 NaCl, 2.5 KCl, 2 CaCl_2_, 2 MgCl_2_, 25 HEPES, and 30 glucose [pH 7.4]) ± isoflurane was delivered using a custom pressurized, inline heated, gas-tight superfusion system [41] at 2 ml/min (equilibration time constant of ∼8s). Cells were equilibrated with control or anesthetic solutions prior to start of experiments and anesthetic concentrations from bath and syringe samples were confirmed by gas chromatography [65]. TrkB-Fc (1μg/ml; R and D Systems, Minneapolis, MN) and recombinant BDNF (75-100ng/ml; PeproTech, Rockyhill, NJ; [18]) were added 5 min prior to stimulation.

### Measurement of exocytosis

Live-cell time-lapse fluorescence images at 10 Hz were collected using a 40x 1.3 (NA) Plan APO oil immersion objective and a Zeiss (Thornwood, NY) Axio Observer 7 microscope equipped with a chamber temperature-controlled at 37°C (Warner Instruments, Hamden, CT). Samples were excited using Colibri 7 solid state LED lamps (469 nm or 555 nm) and fluorescence emission was selected through the 514/30 nm or 592/25 nm band-pass filters (Zeiss, Thornwood, NY). A series of time-lapse images were acquired every 50 ms for 2 min with an Andor iXon 888 electron multiplying CCD camera (Andor, South Windsor, CT) using Zen 2.3 imaging software (Zeiss, Thornwood, NY). The time courses of pHluorin fluorescence responses under tetanic stimulation [2 min; 16 bursts of 50 action potentials (AP) at 50 Hz every 2.5 s] were measured for paired control and treatment conditions in Tyrode’s buffer in the following order: 1) BDNF-pH: 5 min control solution equilibration; 5 s baseline perfusion; 2 min depolarizing tetanic stimulation in control solution; 5 min isoflurane equilibration; 5 s baseline isoflurane perfusion; 2 min depolarizing tetanic stimulation in isoflurane solution; 5 min washout in control solution; 50 mM NH_4_Cl alkalization; 2) vGlut1-pH: 5 min control solution equilibration; 5 s baseline perfusion; 2 min depolarizing tetanic stimulation in control solution; 5 min isoflurane equilibration; 5 s baseline isoflurane perfusion; 2 min depolarizing tetanic stimulation in isoflurane solution; 5 min washout in control solution; 2 min depolarizing tetanic stimulation in control solution; 50 mM NH_4_Cl alkalization. Electrical stimulation was performed using platinum-iridium electrodes and multiple successive time-control stimulations for tetanic pulses were used to rule out fluorescence decay over time.

### Image and statistical analysis

Since secretion of BDNF is asynchronous, we modified our analysis from Shimojo et al. (2015; [66]) as follows: Regions of interest (ROIs) were determined from a change in fluorescence (verified by NH_4_Cl alkalization) where fluorescence intensity is calculated for each ROI and counted as an event if mean Δ F is > 4 SD above baseline. For vGlut1-pH, transfected boutons were selected as ROIs based on their response to NH_4_Cl and/or labeling with mCherry. Each bouton/ROI was subjected to a signal-to-noise calculation based on its response to the first control electrical stimulation, and ΔF was calculated as the difference in average intensities between F_peak_ and F_baseline_ for BDNF-pH and ΔF/F calculated as the difference in average intensities between F_pea_k and F_baseline_/F_baseline_ for vGlut1-pH. Release probability were analyzed as mean number of fusion events (BDNF; [66]) or mean (vGlut1; [67]) ΔF/F normalized to the total vesicle pool determined by alkalization with 50 mM NH_4_Cl to compensate for differences in density of available vesicles across independent imaging experiments. Selection of ∼30-50 ROIs from each neuron included both distal and proximal regions relative to the soma for BDNF-pH and axonal boutons for vGlut1-pH. Fluorescence data were analyzed using Time Series Analyzer (ImageJ) for vGlut-pH or FluoroSNAP (Matlab) for BDNF-pH. Data are expressed as mean ± SD. Drug effects are shown as a percentage of total pool release or control response to allow quantification of inhibition or potentiation. Statistical significance was determined by unpaired two-tailed Student’s *t* tests or one-way ANOVA with Tukey’s *post hoc* test, with *p* < 0.05 considered significant. Statistical analysis and graph preparation were performed using GraphPad Prism v7.05 (Graphpad Software, Inc., San Diego, CA).

## Results

### Isoflurane reversibly inhibits excitatory synaptic vesicle exocytosis

Reductions in SV exocytosis by general anesthetics vary with type of neuron [33, 37] and can be mechanistically distinct based on stimulation frequency [33, 47]; with short depolarizing pulses, anesthetics reduce Ca^2+^ influx independent of Ca^2+^-exocytosis coupling [33, 47], while with longer depolarizations anesthetics directly inhibit the SV exocytotic machinery downstream of Ca^2+^ influx [47]. With short physiological stimulations, isoflurane transiently inhibits transmitter release from hippocampal glutamatergic and GABAergic or ventral tegmental area (VTA) dopaminergic neurons [33, 37]. Patterns of both high and low brain activity are observed under anesthesia in both rodents and humans [68, 69], but little research has focused on long, repetitive input, particularly in the hippocampus, an area critical for synaptic plasticity and BDNF signaling.

Rat hippocampal neurons were exposed to isoflurane (0.53-0.7 mM; ∼1.5-2 MAC) and stimulated with high-frequency tetanic pulses (16 bursts of 50 action potentials at 50 Hz every 2.5 s [70, 71] to examine the effects of isoflurane on vGlut-pH SV exocytosis (Fig. 1A; supplemental movie S1). Presynaptic boutons were visualized using mCherry (Fig. 1B) and exocytosis at glutamatergic nerve terminals was measured by the fluorescence change in pH-sensitive pHluorin fused to the luminal domain of the vesicular glutamate transporter (vGlut-pH; [67]; Fig.1C). Fluorescence responses were normalized to total SV pool by NH_4_Cl alkalization and to control for between neuron variability. The vGlut-pH biosensor reliably measured SV exocytosis over time with minimal decay over the course of three control stimulations (data not shown). In glutamatergic neurons, stimulated exocytosis was 64% of total pool (TP), which was reduced to 44% of TP by 0.7 mM isoflurane (31% inhibition), a clinically relevant concentration ∼ 1.4 times the EC_50_ for general anesthesia in rat [64]. Exocytosis returned to control levels (72% of TP) within 3 min of isoflurane wash-out under the same stimulation conditions (Fig. 1A, D). This recovery in SV release was stable unless cell health was compromised due to photobleaching or toxicity. These results show that compared to lower depolarization frequencies [33, 37], release as a fraction of total pool is greater with high frequency stimulation, but is comparably reversible and recovers following isoflurane washout.

**Figure 1.**
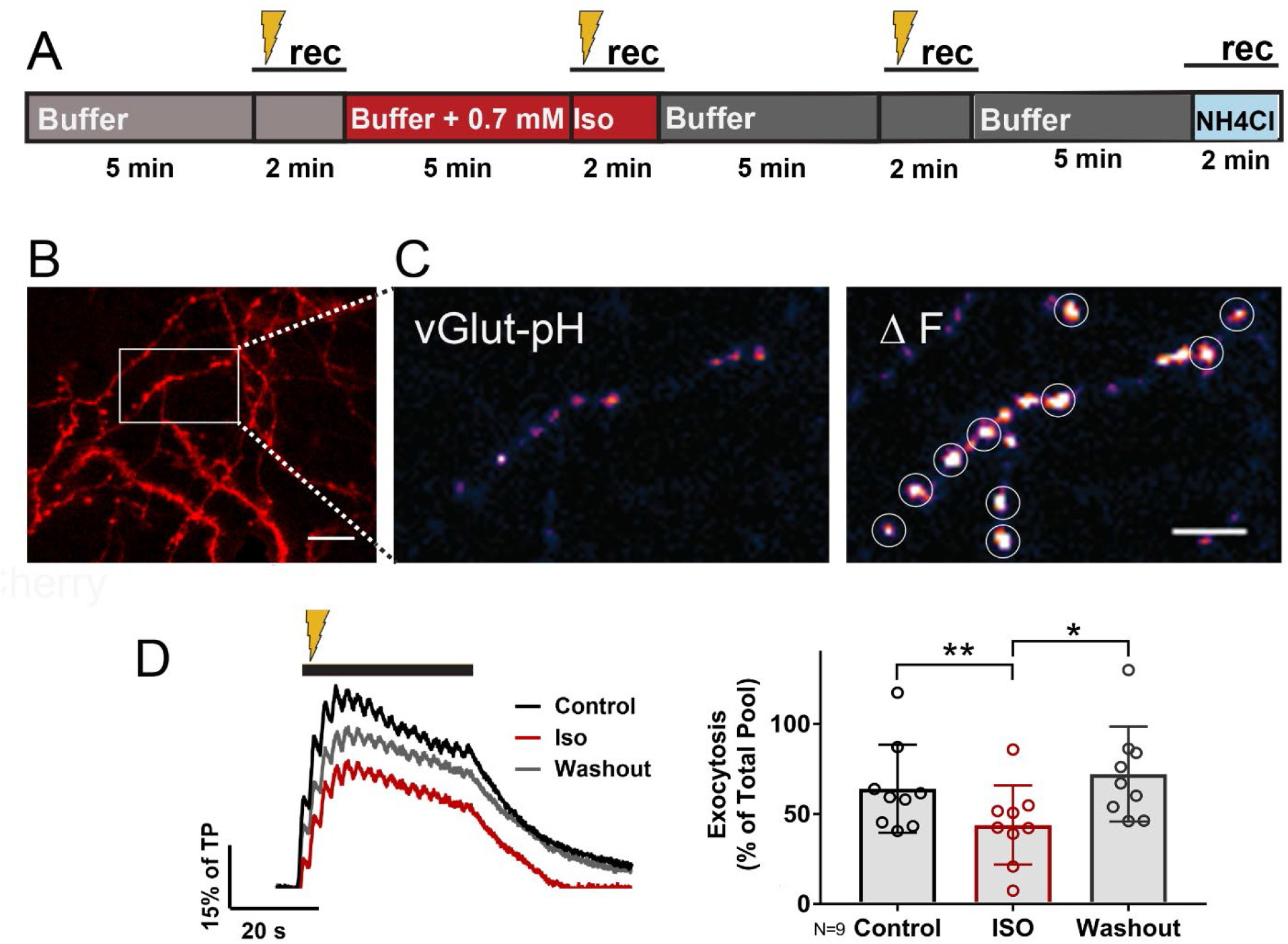
Isoflurane reversibly inhibits electrically-evoked synaptic vesicle exocytosis. Rat hippocampal neuron cultures (16DIV) transfected with vGlut-pH and mCherry were perfused with 0.5-0.7 mM isoflurane or control buffer with tetanic stimulation (gold lightning bolt) as indicated in **(A)**. Representative images showing dendritic arbor and axons of a control neuron (**B**-mCherry). Inset shows single ROIs of boutons (vGlut-pH) before (**C**-left) and after (**C**-right, white circles) tetanic stimulation. Electrically-evoked synaptic vesicle exocytosis (**D**-left, gold bar) was reduced by isoflurane **(D**-right) (**p<0.01 by one-way multiple comparisons ANOVA with Dunnett’s *post hoc* test). Reduced exocytosis observed with isoflurane recovered following washout (**D**-right) (*p<0.05 by one-way multiple comparisons ANOVA with Dunnett’s *post hoc* test). Data are mean ± SD %; n=9 neurons per experimental group and 405-450 boutons. Scale bar = 5 μm.

### BDNF contributes to excitatory synaptic vesicle exocytosis

In the mature brain, mBDNF is a key modulator of synaptic plasticity that enhances glutamate release by multiple mechanisms: activation of phospholipase Cγ (PLCγ) increases intracellular Ca^2+^ [55]; phosphorylation of synapsin by mitogen-activated protein kinase (MAPK) [56] and increased Rab3 expression [57] enhance exocytosis; and activation of the actin motor complex increases neurotransmitter replenishment and release [58, 59, 72]. We tested the contribution of BDNF to SV exocytosis under high-frequency tetanic pulses (16 bursts of 50 action potentials at 50 Hz every 2.5 s) by using TrkB-Fc (1 μg/ml), a chimeric protein containing the extracellular domain of TrkB that binds and sequesters endogenous BDNF. Reduction of extracellular BDNF levels by TrkB-Fc inhibited tetanic stimulation-evoked SV exocytosis by 17% as quantified using vGlut-pH in rat hippocampal neurons (Fig. 2A). Quantification of vGlut-pH without electrical stimulation confirmed that exogenous addition of mBDNF (75-100 ng/ml) significantly increased basal SV exocytosis (Fig. 2B,C). Compared to the modest reduction of exocytosis with TrkB-Fc with tetanic stimulation, the large increase in vGlut-pH fluorescence observed with exogenous mBDNF is likely due losses from peptide delivery. Based on the role of BDNF on neurotransmitter release, we posit that BDNF may be a candidate target for anesthetic-induced effects on synaptic transmission.

**Figure 2.**
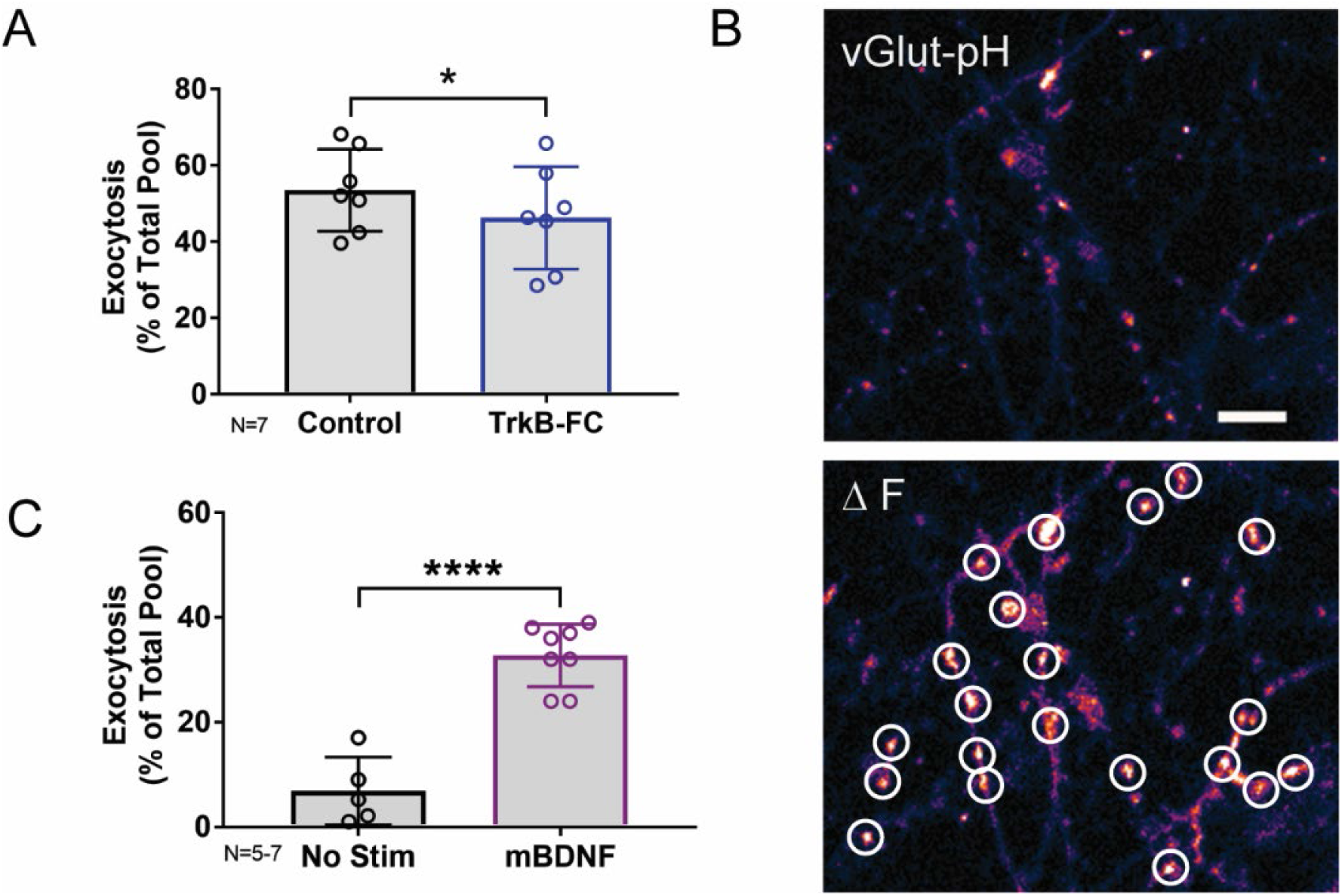
Endogenous BDNF contributes to synaptic vesicle exocytosis. Quantification of synaptic vesicle exocytosis shows that pretreatment with the BDNF binding protein TrkB-Fc (1 μg/ml) for 5 min prior to tetanic stimulation reduced exocytosis **(A)** (*p<0.05 by Student unpaired t-test). Representative images of a rat hippocampal neurons (16DIV) transfected with mCherry (not shown) and vGlut-pH showing single ROIs of boutons before (**B**-top) and after (**B**-bottom, white circles) addition of exogenous mBDNF (75-100 ng/ml), which increased synaptic vesicle exocytosis (**C**) (****p<0.0001 by Student unpaired t-test). Data are mean ± SD; n=5-7 neurons per experimental group and 250-350 boutons. Scale bar = 5 μm.

### Attenuation of BDNF release by isoflurane contributes to depression of excitatory synaptic vesicle exocytosis

Multiple presynaptic targets have been identified for volatile anesthetic actions [39-41, 73, 74], but they do not fully explain anesthetic-induced reduction in excitatory transmission. As BDNF modulates glutamate release in an isoflurane-sensitive manner (Fig. 2), we tested the role of BDNF in the effects of isoflurane by determining isoflurane-induced changes in activity-dependent BDNF release in live hippocampal cells using pHluorin fused to the C-terminus of BDNF (BDNF-pH). BDNF release can be induced electrically by high-frequency tetanic stimulation and BDNF-pH is properly processed, asynchronously released (Fig. 3 A,B; supplemental movie S2), and biologically active following secretion [71]. Exocytosis of BDNF was inhibited ∼47% by isoflurane (Fig. 3C) suggesting a novel role in contributing to isoflurane-induced effects of synaptic transmission. Taken together, our data show that BDNF contributes to SV exocytosis (Fig. 2) and isoflurane inhibits BDNF release (Fig. 3). To delineate the consequence of reduced BDNF release by isoflurane on glutamatergic exocytosis, we examined isoflurane-induced reduction of SV exocytosis using vGlut1-pH in the presence of TrkB-Fc to sequester extracellular BDNF. Reducing extracellular BDNF attenuated depression of excitatory SV exocytosis by isoflurane (Fig. 4) showing contribution of BDNF signaling to modulation of excitatory transmission by isoflurane.

**Figure 3.**
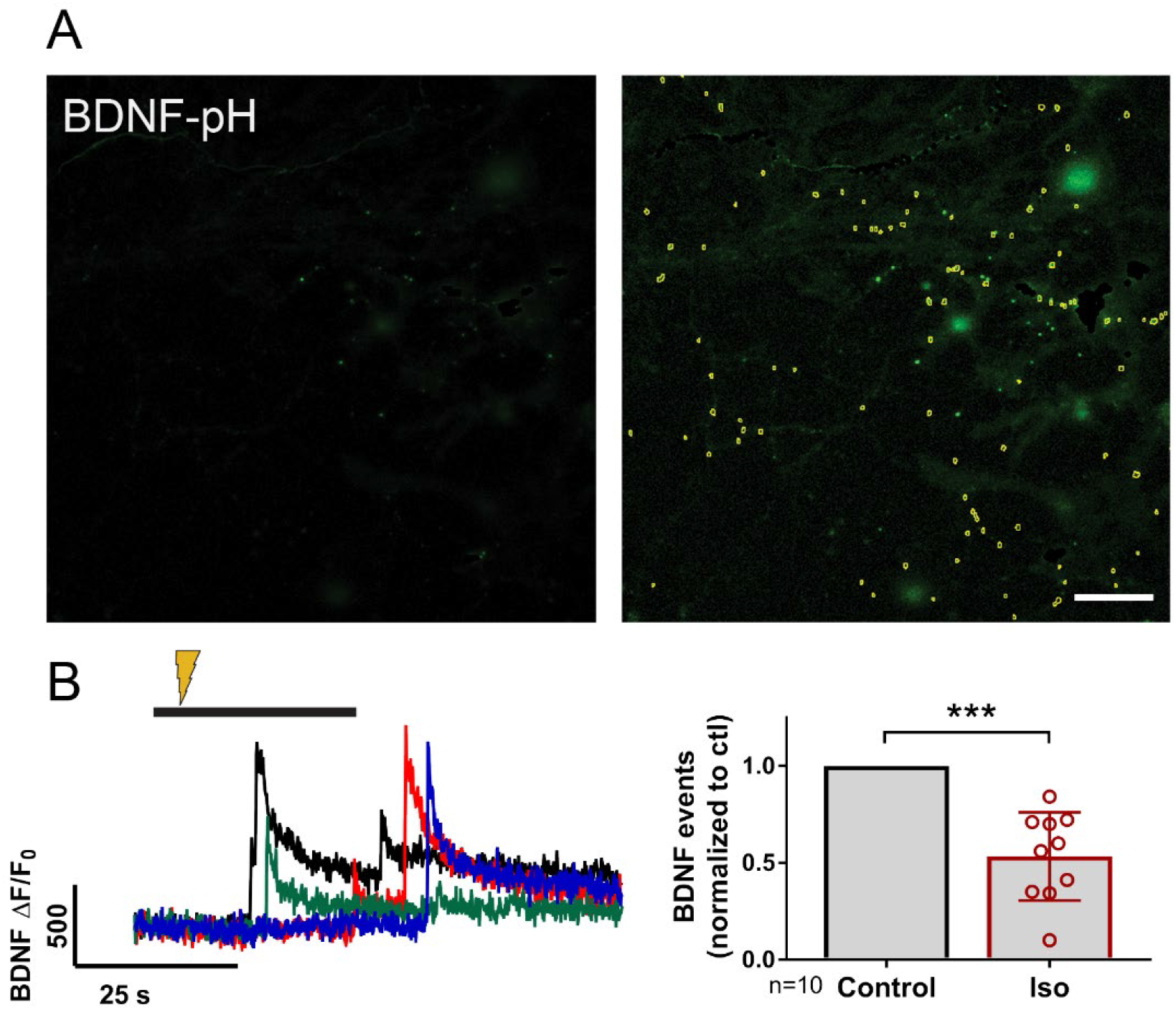
BDNF exocytosis is reduced by isoflurane. Representative images of rat hippocampal neurons (18DIV) transfected with mCherry (not shown) and BDNF-pH showing single BDNF exocytotic events (yellow dots) before **(A-**left) and after (**A**-right) tetanic stimulation (**B**-left, gold lightning bolt) showing asynchronous release of BDNF-pH (**B**-left). Quantification of BDNF exocytosis defined using fluorescence responses to tetanic stimulation showed inhibition by 0.5-0.7 mM isoflurane (**B**-right) (***p<0.001 by Student unpaired t-test). Data are mean ± SD; n=10 neurons per experimental group and 23-113 BDNF events selected for each condition. Scale bar = 10 μm.

**Figure 4.**
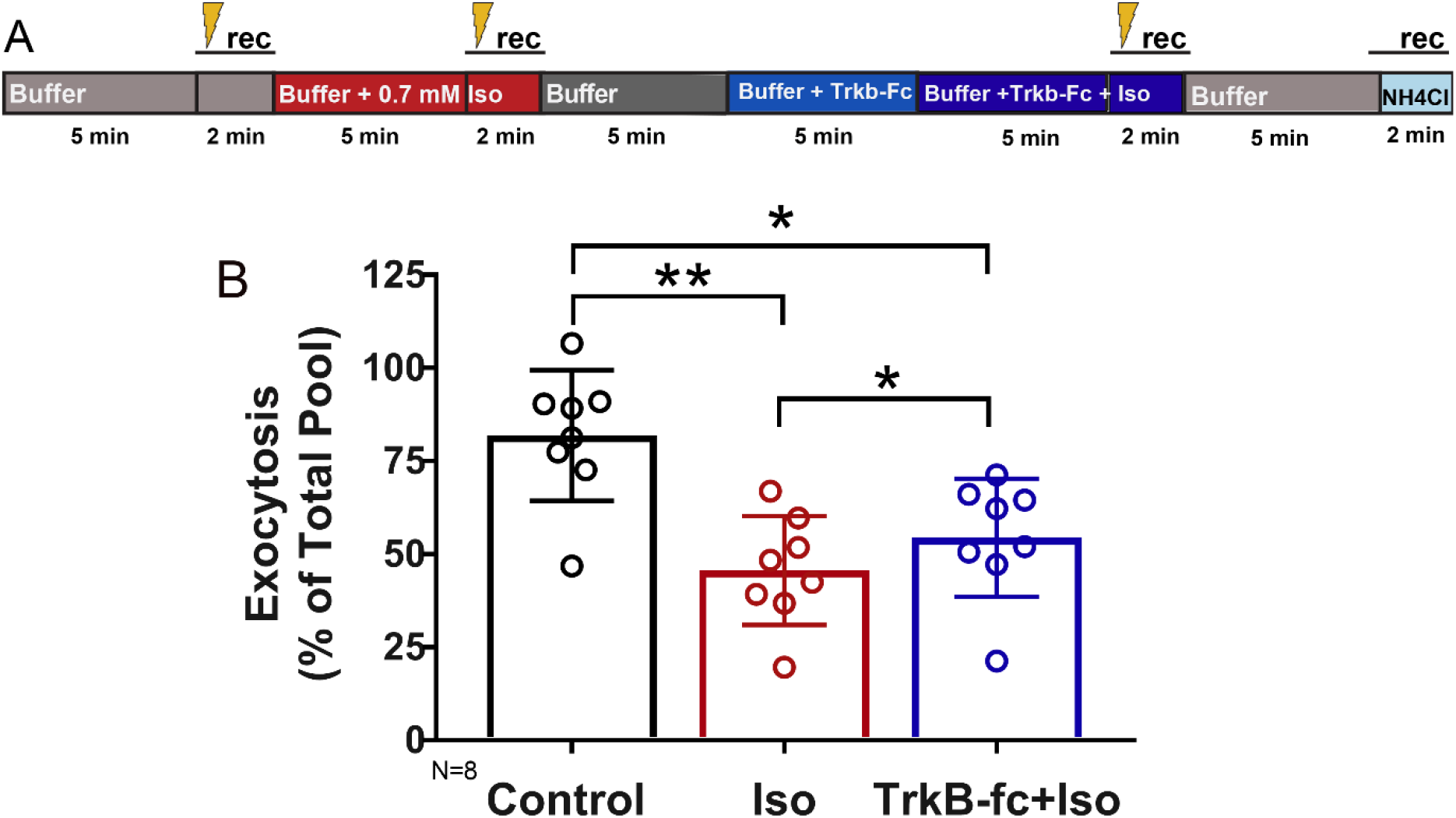
Inhibition of synaptic vesicle exocytosis by isoflurane is attenuated by reducing endogenous BDNF. Rat hippocampal neuron cultures (16DIV) transfected with vGlut-pH and mCherry were perfused with control buffer, 0.5-0.7 mM isoflurane, or TrkB-fc + 0.5-0.7 mM isoflurane with tetanic stimulation (gold lightning bolt) as indicated in **(A). (B)** Synaptic vesicle exocytosis evoked by tetanic stimulation was reduced by isoflurane (0.5-0.7 mM) (**p<0.01 by multiple comparisons one-way ANOVA with Dunnett’s *post hoc* test). Reduced exocytosis observed with isoflurane exposure was attenuated by pretreatment with TrkB-Fc (1 μg/ml) for 5 min prior to stimulation (*p<0.05 by multiple comparisons one-way ANOVA with Dunnett’s *post hoc* test). Data are mean ± SD; n=8 neurons per experimental group and 327-400 boutons.

### Neurons expressing Val66Met and Met66Met BDNF show sustained depression of SV exocytosis by isoflurane compared to Val66Val neurons

Attenuation of excitatory transmission leads to morphological changes in hippocampal dendritic spines [75], and persistent changes in spine structure are associated with cognitive dysfunction in various neurological disorders [76]. BDNF regulates spine size and knock-in mice with the BDNF SNP (BDNF Val66Met) have altered spine morphology and density in the medial prefrontal cortex, basolateral amygdala [77] and hippocampus [78]. However, alterations in presynaptic structure and function due to the Val66 genotype have not been investigated. The Val66Met SNP results in abnormal intracellular trafficking of BDNF leading to lower regulated secretion of mBDNF [23-25], but did not directly influence glutamate release or total amount of releasable pool (Fig. 5B) as assayed by vGlut1-pH with tetanic stimulation. Presynaptic bouton size and number are difficult to determine in unpaired cells due to heterogeneity of the cell population in culture and architecture of the axon, but qualitative differences were not found between Val66Met and Met66Met boutons compared to Val66Val wild-type boutons (Fig. 5A).

**Figure 5.**
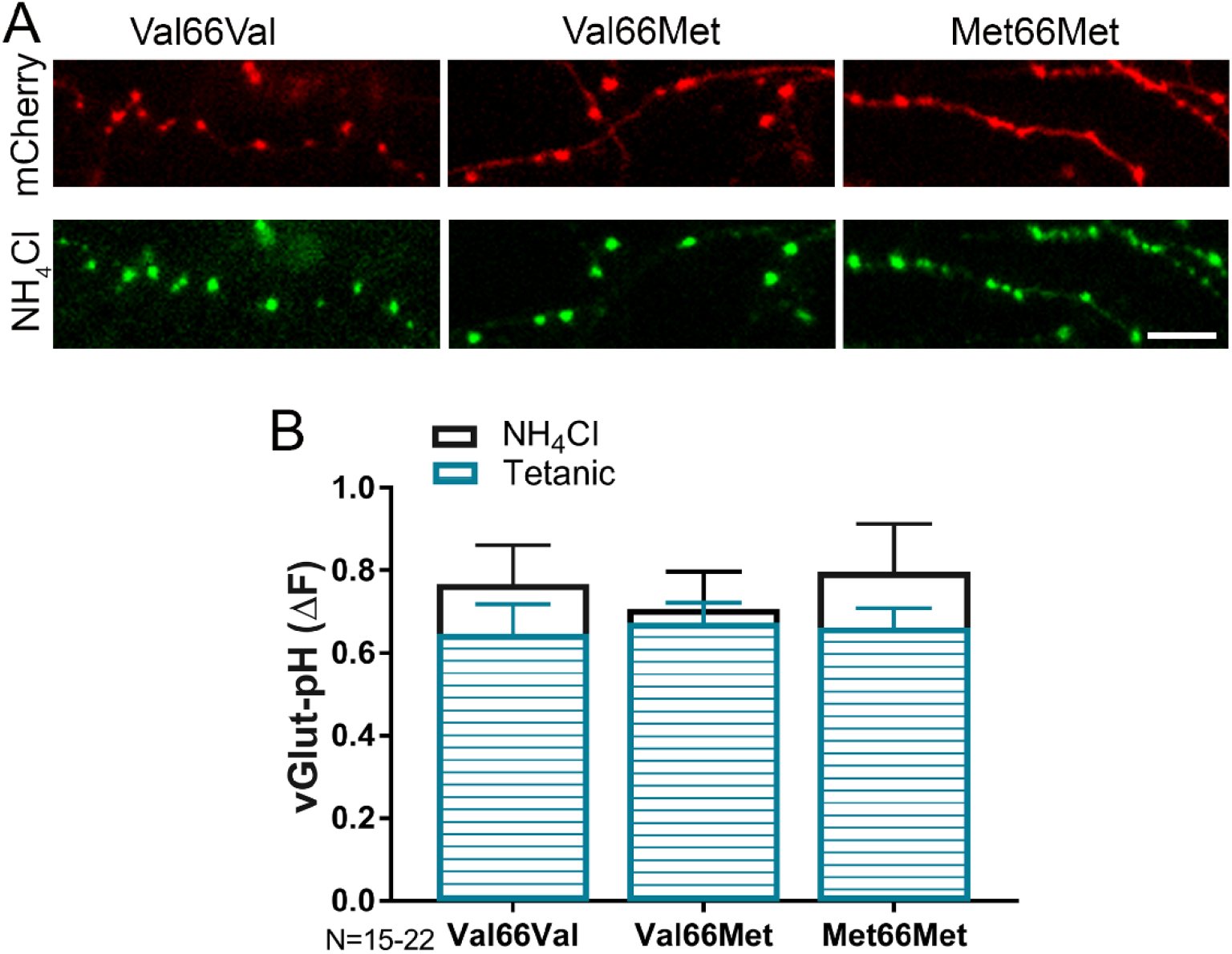
BDNF Val66Met and Met66Met single nucleotide polymorphisms do not affect total synaptic vesicle number or amount of exocytosis. Representative images of 14-16DIV Val66Val (wt), Val66Met, and Met66Met mouse hippocampal neurons transfected with mCherry (**A**-top) and vGlut-pH (with NH_4_Cl alkalization, **A**-bottom). (**B**) Change in fluorescence of vGlut-pH with NH_4_Cl alkalization or tetanic stimulation reflecting total vesicle number (**B**-clear) or amount of exocytosis (**B**-striped) showed no significant differences between the three genotypes. Data are mean ± SD; n=15-22 neurons per experimental group and 750-1100 boutons. Scale bar = 5 μm.

Since BDNF is a target for isoflurane-induced reduction of synaptic transmission (Fig. 4), neurons with reduced BDNF levels were less affected by isoflurane exposure (Fig.6A; % inhibition: wt-44%; Val66Met-22%; Met66Met-30%). However, inhibition of SV exocytosis by isoflurane was irreversible in Val66Met and Met66Met neurons compared to Val66Val (Fig. 6A) following drug washout. These changes were rescued by addition of exogenous mBDNF (Fig. 6B), confirming the contribution of the BDNF SNP in producing prolonged structural and functional presynaptic changes with consequences for isoflurane anesthesia.

**Figure 6.**
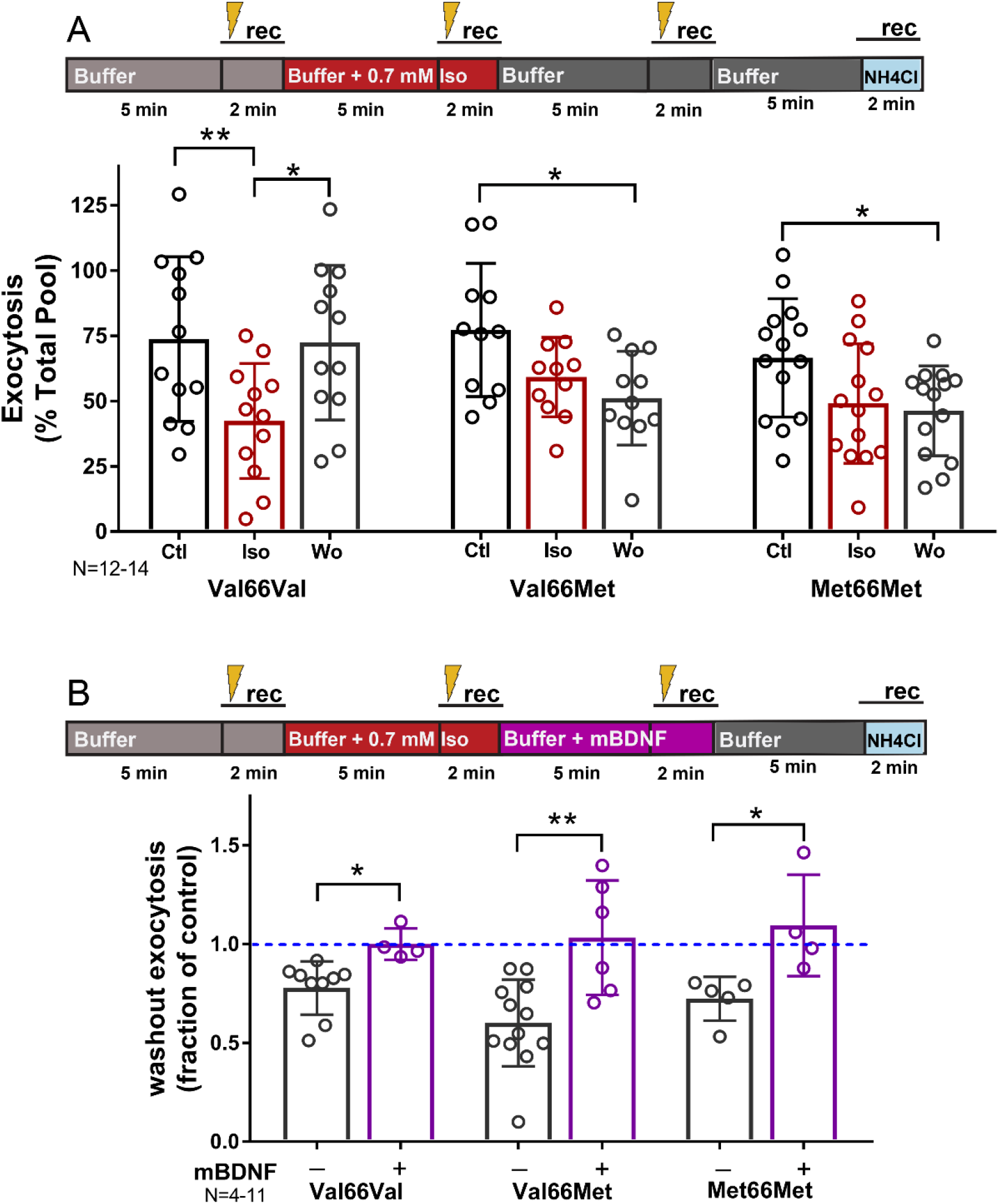
Inhibition of synaptic vesicle exocytosis by isoflurane is irreversible in Val66Met and Met66Met neurons, but rescued with exogenous BDNF. Mouse hippocampal neuron cultures (16DIV) transfected with vGlut-pH and mCherry were perfused with control buffer, 0.5-0.7 mM isoflurane, or mBDNF with tetanic stimulation (gold lightning bolt) as indicated in **(A, B**, top). Tetanic stimulation of synaptic vesicle exocytosis measured with vGlut-pH was inhibited by isoflurane (0.5-0.7 mM) in Val66Val mouse neurons compared to Val66Met and Met66Met neurons (**p<0.01 by multiple comparisons one-way ANOVA with Dunnett’s *post hoc* test). Changes in vesicle exocytosis observed in Val66Met and Met66Met neurons after isoflurane exposure did not recover following washout of drug (**A**, bottom) (*p<0.05 by multiple comparisons one-way ANOVA with Dunnett’s *post hoc* test). Addition of 75-100 ng/ml of mBDNF during washout rescued synaptic vesicle exocytosis in Val66Met and Met66Met and increased exocytosis in Val66Val (**B**, bottom) (*p<0.05; **p<0.01 neurons by Student unpaired t-test). Data are mean ± SD; n=4-14 neurons per experimental group and 200-700 boutons.

## Discussion

We found that isoflurane inhibits glutamatergic SV exocytosis evoked by high frequency stimulation in part by reducing release of endogenous neuromodulator BDNF. Moreover, neurons with reduced endogenous BDNF levels as a result of BDNF SNPs show persistent reductions in glutamatergic exocytosis following exposure to isoflurane (Fig.7). Isoflurane effects on BDNF signaling provide a novel mechanism for its acute neurophysiological effects that may contribute to lasting impairments in synaptic plasticity and cognitive function based on genotype.

**Figure 7.**
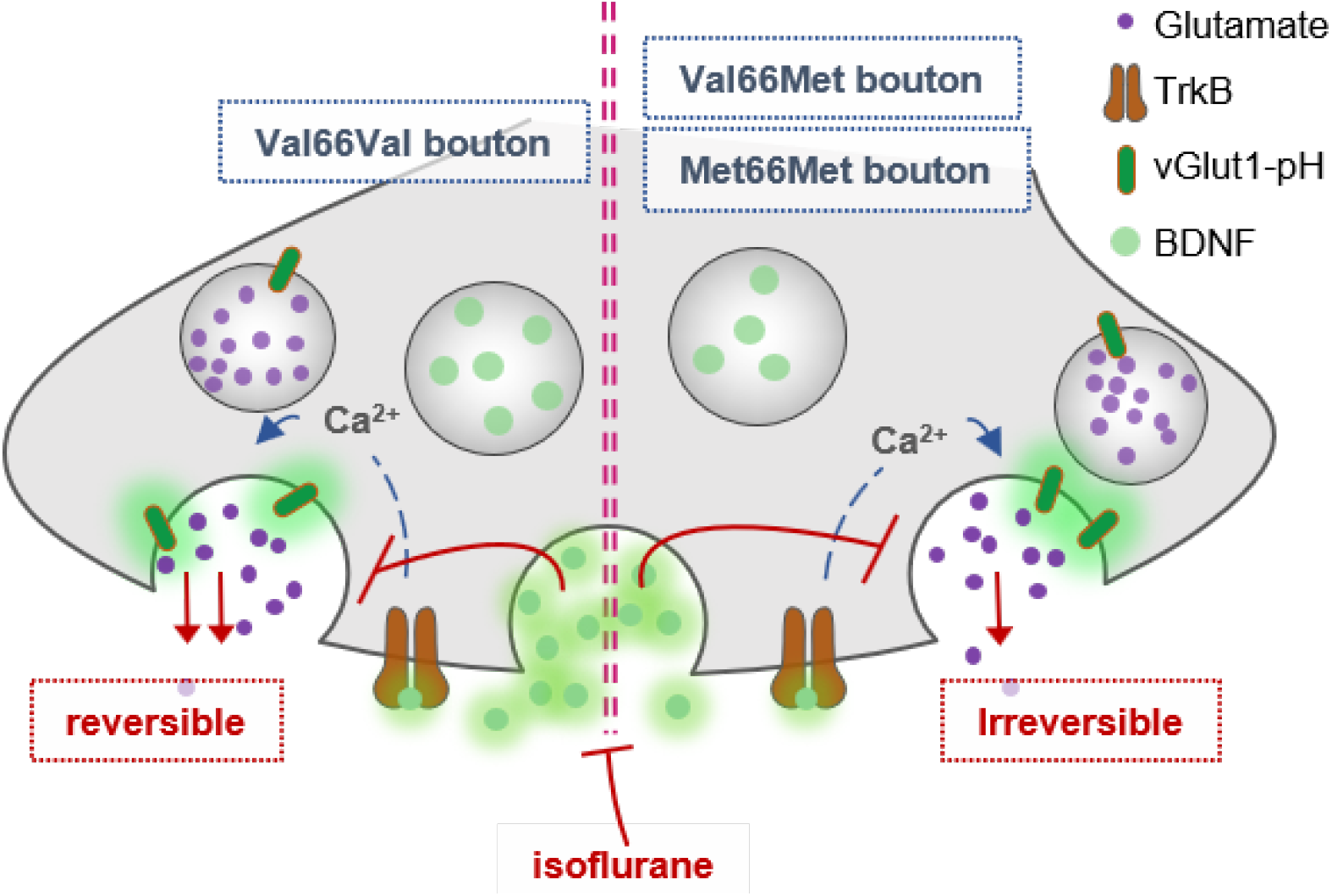
Role of BDNF in isoflurane-induced depression of synaptic transmission. BDNF signaling contributes to glutamatergic synaptic vesicle exocytosis. Isoflurane inhibits glutamatergic synaptic vesicle exocytosis evoked by tetanic stimulation in part by reducing release of endogenous BDNF. Neurons with reduced endogenous BDNF levels as a result of BDNF single nucleotide polymorphisms (Val66Met; Met66Met) show persistent reductions in glutamatergic exocytosis following exposure to isoflurane.

Our findings confirm previous studies [55, 56, 58] that showed that activity-dependent BDNF signaling influences excitatory synaptic transmission by enhancing glutamate exocytosis. We now report that these mechanisms are relevant to isoflurane effects on synaptic function since BDNF release evoked by tetanic stimulation was significantly inhibited by clinically relevant concentrations of isoflurane and contributed to its depression of excitatory SV exocytosis. Previous studies have implicated proBDNF in anesthetic neurotoxicity in developing neurons and astrocytes [79-83], but did not investigate the role of mBDNF in established hippocampal cultures. Head and colleagues [80] showed that isoflurane produces neurotoxicity through effects on proBDNF signaling mediated by p75 receptors that promote cell death and attenuate synaptic transmission [84, 85]. Upregulating cleavage of proBDNF to mBDNF or direct inhibition of p75 receptors reduces both anesthetic-induced neuronal apoptosis [80] and destabilization of dendritic filopodia in developing neurons [79], but the direct effects on mBDNF were not reported. Our findings suggests that novel mBDNF mechanisms are involved in the acute and transient effects of isoflurane in reduction of excitatory transmission in established hippocampal boutons.

We found that with high-frequency tetanic stimulation, isoflurane reversibly inhibited excitatory SV exocytosis in wild-type neurons consistent with transient modification of synaptic function. This is consistent with previous studies showing no persistent changes in depolarization-evoked neurotransmitter release for mechanistically distinct anesthetics [33, 37, 86], although distinct mechanisms are posited based on stimulation [47]. Excitatory transmission is critical to hippocampal synaptic plasticity, and can be modulated by anesthetics to produce long-term neurocognitive consequences [87, 88]. Genetic variation is a risk factor for postoperative neurocognitive decline [10], and the BDNF Val66Met allele is a polymorphism associated with susceptibility to postoperative neurocognitive decline [8, 26, 27]. This provides a mechanism for long-term changes in synaptic connectivity and function that persist beyond acute anesthetic actions, possibly contributing to cognitive dysfunction.

The biochemical, anatomical, and behavioral consequences of the human BDNF Val66Met polymorphism *in vivo* are recapitulated in a Val66Met BDNF knock-in mouse model [89, 90]. These mice have attenuated long-term potentiation (LTP; [91]) and reduced hippocampal neuronal soma size and dendritic complexity [90]. Regulated BDNF secretion is reduced by 18% in Val66Met and 30% in Met66Met neurons [90] compared to wild-type, allowing comparisons of ‘dose-dependent’ BDNF effects in disruption of synaptic function by isoflurane. We found that in Val66Met and Met66Met mouse hippocampal boutons isoflurane induced sustained depression of SV exocytosis compared to Val66Val boutons. This suggests that although the contribution of mBDNF to glutamate release is modest (Fig. 2), it is a critical component of synaptic function in the presence of isoflurane. Whether these alterations in synaptic function are accompanied by changes in synaptic structure and connectivity will require further study. Consistent with a potential role of regulated BDNF release in synaptic transmission, the Met66Met genotype impairs NMDA receptor-mediated synaptic transmission and plasticity compared to wild-type in the hippocampus [91]. Reduced synaptic transmission by the BDNF SNP is region-dependent, as enhanced glutamatergic transmission was revealed in the dorsolateral striatum of BDNF Met66Met mice compared to wild-type, but functionally still resulted in impairments of LTP and LTD [92].

Our results did not find genotypic differences between Val66Val, Val66Met, or Met66Met boutons in SV total pool size or vGlut1-pH exocytosis with tetanic stimulation, implicating a role for reduced mBDNF in modulating the persistent presynaptic changes by isoflurane. This BDNF-dependent effect on the irreversibility of isoflurane inhibition of SV exocytosis in Val66Met and Met66Met boutons was verified by rescue with exogenous mBDNF (Fig. 6). The magnitude of SV exocytosis with mBDNF was higher than that of control stimulations suggesting that basal transmission was sub-maximally stimulated by endogenous mBDNF. Exogenous mBDNF can induce hippocampal plasticity both in *in vivo* [93] and in *in vitro*, with short-term synaptic enhancement occurring at 20 ng/ml *in vitro* [94]. However, we did not observe an effect below 75 ng/ml most likely due to losses during perfusion. The intersection of BDNF SNPs with isoflurane-induced changes in glutamatergic transmission and plasticity should be confirmed with electrophysiological measurements.

BDNF is a secreted growth factor packaged in and released form large dense-core vesicles. It is constitutively released but can also be released in mature neurons by prolonged depolarization, high-frequency, theta-burst, or tetanic stimulations [95, 96] which have physiological correlates in synaptic transmission and plasticity [71, 97]. We studied anesthetic effects on BDNF release in real-time using the fluorescent fusion protein BDNF-pH, which can be released by tetanic stimulation [70, 71]. The fluorescence of BDNF-pH is quenched by the acidic interior of dense core vesicles where BDNF is stored for release and is rapidly unquenched when these vesicles fuse with the plasma membrane. BDNF can be released from axons or dendrites [71, 98] and has diverse pre- and post-synaptic modulatory effects on mature glutamatergic synapses consistent with our evidence as a presynaptic target for anesthetics. Using BDNF-pH, Matsuda et al. examined stimulation-evoked BDNF secretion from dissociated rat hippocampal neurons. With brief stimuli, BDNF secretion was predominantly at dendrites compared to axons, while prolonged high-frequency stimulation was necessary for vesicular fusion from axons [71]. Based on our stimulation paradigm, these findings support anesthetic-mediated effects on axonal BDNF, but further studies are needed to delineate possible site-specific actions. To explore all possible sites of BDNF release, we did not include blockers of postsynaptic NMDA or AMPA receptors in our protocols as BDNF release from dendritic spines is largely NMDA-dependent [99]. However, we performed time control experiments with three successive stimulations to exclude postsynaptic potentiation and to ensure fluorophore stability (data not shown).

The mechanisms for acute depression of excitatory synaptic transmission by general anesthetics include reduction of neuronal excitability [28] or action potential conduction [29-32], inhibition of presynaptic Ca^2+^ influx [33, 34] and synaptic vesicle (SV) exocytosis [35-37], and/or blockade of postsynaptic glutamate receptors [38]. Based on our current findings, further studies should examine the contribution of BDNF and the role of genotype to these mechanisms. We have identified a role for reduced BDNF signaling resulting from the Val66Met SNP, which persistently alters synaptic function following exposure to isoflurane. Dissociated hippocampal neurons replicate most of the fundamental cellular and molecular features of synaptic structure and function so our findings are of potential fundamental importance *in vivo*. However, the irreversible deficit in glutamatergic synaptic function produced by isoflurane in the BDNF Met SNPs should be further verified in functional and behavioral assays *in vivo*. Previous studies have observed genetic variation in vulnerability to postoperative neurocognitive dysfunction. Based on our findings additional studies are needed to confirm whether the effects observed in rodent hippocampal neurons can be extrapolated to mechanistically distinct anesthetics, other brain regions, and other neuronal subtypes, and whether they translate to the clinical setting.

## Funding

This work was supported by National Institutes of Health (NIH) grants GM130722 to JP and GM058055 to HCH.

## Supporting Information

**Movie S1. Tetanic stimulation releases glutamatergic synaptic vesicles**. Rat hippocampal neuron cultures (16DIV) transfected with vGlut1-pH and mCherry (not shown) were perfused with Tyrode’s control buffer with tetanic stimulation (16 bursts of 50 action potentials at 50 Hz every 2.5 sec).

**Movie S2. Tetanic stimulation releases BDNF**. Rat hippocampal neuron cultures (18DIV) transfected with BDNF-pH and mCherry (not shown) were perfused with Tyrode’s control buffer with tetanic stimulation (16 bursts of 50 action potentials at 50 Hz every 2.5 sec).

## Notes

### Competing Interest Statement

The authors have declared no competing interest.

